# Covering all your bases: incorporating intron signal from RNA-seq data

**DOI:** 10.1101/352823

**Authors:** Stuart Lee, Albert Y. Zhang, Shian Su, Ashley P. Ng, Aliaksei Z. Holik, Marie-Liesse Asselin-Labat, Matthew E. Ritchie, Charity W. Law

## Abstract

RNA-seq datasets can contain millions of intron reads per sequenced library that are typically removed from downstream analysis. Only reads overlapping annotated exons are considered to be informative since mature mRNA is assumed to be the major component sequenced, especially when examining poly(A) RNA samples. In this paper, we demonstrate that intron reads are informative and that pre-mRNA is the major source of intron signal. Making use of pre-mRNA signal, our *index* method combines differential expression analyses from intron and exon counts to categorise changes observed in each count set, giving additional genes with evidence of transcriptional changes when compared to a classic approach. Considering the importance of intron retention in some biological systems, another novel method, *superintronic*, looks for evidence of intron retention after accounting for the presence of pre-mRNA signal. The results presented here overcomes deficiencies and biases in previous works related to intron reads by exploring multiple sources for intron reads simultaneously using a data-driven approach, and provides a broad overview into how intron reads can be utilised in relation to multiple aspects of transcriptional biology.

## Introduction

Advances in gene profiling technology, such as RNA-sequencing (RNA-seq) have allowed researchers to study transcription in exquisite detail. Previously, quantitative gene expression analyses by microarray required prior knowledge of the sequences to be interrogated, limiting *de novo* discoveries and understanding of gene transcripts and alternative splicing especially at a high-throughput level. Most research efforts focused on gene-level information and comparison of genes that are differentially expressed (DE) between two or more groups. Whilst this is still the main focus for RNA-seq, the technology has the ability to examine sub-gene components such as at the transcript-level, exon-level, or even nucleotide base-level without prior sequence knowledge. As a result, there has been increased interest and effort into the study of transcript-level information, alternative gene splicing and gene intron retention (IR) at a global level using RNA-seq (1–6).

RNA-seq can be used to characterise and study many RNA types, including non-coding RNAs that regulate a diverse range of cellular processes (7, 8), but the overwhelming majority of studies focus on messenger RNAs (mRNAs) which encode genes that are translated into protein. The most popular RNA selection protocol is that which captures polyadenylated RNA (poly(A) RNA) seeing that it is optimised for mRNA selection – in other words, RNA that has a poly(A) tail at the 3’ end of the molecule. In eukaryotes, poly(A) tails are synthesised to aid in the transportation of mature mRNA molecules from the nucleus to the cytoplasm. Total RNA selection is also widely used and often includes a step that depletes ribosomal RNA so that it does not compete with sequencing of mRNA. RNA expression values are highly correlated between the two RNA selection protocols, with a higher percentage of reads (≈3% more) mapping to protein coding genes in poly(A) RNA samples, and a higher percentage of reads (≈2.5% more) mapping to long non-coding RNAs in Total RNA samples (9). The general assumption is that for protein coding genes the vast majority of RNA captured during the RNA-seq experiment are mature mRNA transcripts – this is especially true when it comes to experiments for poly(A) RNA samples. As a result, aligned sequencing reads are usually summarised only for annotated exons within genes. However, it has been shown that intron reads account for a significant proportion of sequencing reads (10–12) yet it is not common practice to quantify these reads. Perhaps this is due to suggestions that intron reads represent experimental and transcriptional noise (13), or are unusable in exon and gene quantification (14).

Other studies have demonstrated that intron read changes are correlated with measurements of nascent RNA (15) or have used intron reads to detect genes with retained introns (4, 16), where IR has been shown to play important roles in inactivation of tumour suppressor genes (17) and regulation of gene expression during neutrophil (18) and erythroblast (19) differentiation. In the former scenario where Gaidatzis *et al.* (2015) (15) studied nascent transcription in both poly(A) RNA and Total RNA libraries, IR is not mentioned as a possible source of intron reads. In the latter, Wong *et al.* (2013) (16) assumed that poly(A) RNA libraries contain processed mRNAs – they show that genes with differentially retained introns are enriched in the cytoplasm.

The literature on the usability of intron reads is ambiguous; it suggests that either intron reads are indicative of nascent transcription or they correspond to retained introns in mature mRNAs. Should an analyst run Gaidatzis’s exon-intron split method (*EISA*) method (15) to separate genes that are transcriptionally regulated to those that are post-transcriptionally regulated between two conditions? Or should they use Wong’s *IRFinder* or like methods (16, 20–23) to examine IR? In practice, the decision could probably be driven by the biology of interest. However, what does this mean for the interpretation of results and are they in contradiction with each other if results are overlapping?

In this paper, the usefulness of intron reads, and the two main competing explanations for observing intron reads are considered in poly(A) RNA and Total RNA libraries. A data-driven approach is taken to understand characteristics of intron and exon reads common to datasets. Technical properties from the data were examined to determine possibility of intron reads resulting from pre-mRNA or IR, or both. In depth biological mechanisms are not discussed as it is beyond the scope of this paper. We demonstrate that intron reads are informative of biological signal and their characteristics indicate that intron reads are predominantly from pre-mRNA rather than IR. Given that both biological processes are important, two novel methods are presented – *index* incorporates intron reads into differential gene expression (DGE) analyses to catergorise changes by transcriptional stage; and *superintronic* examines coverage patterns to select genes with IR-like profiles whilst accounting for underlying pre-mRNA signal. The *index* method was applied to human cell lines and immune cells, showing a unique set of genes where transcriptional changes can be detected in intron counts but not in exon counts, whilst transcriptional changes are mostly consistent between intron and exon counts. *Superintronic* successfully detected genes with IR-like coverage profiles, unlike that of *IRFinder* (20) and *IsoformSwitchAnalyzeR* (23) which appeared to detect genes with pre-mRNA signal and changes in intron coverage but no obvious retained intron was observable from coverage profiles. Allowing for the co-existence of both sources, our study provides an opportunity to both naive and experienced researchers to examine RNA-seq reads more efficiently and for further exploration into key biological processes.

## Materials and Methods

### Datasets

*Human cell lines* of lung adenocarcinoma HCC827 and NCI-H1975 were cultured on three separate occasions by Holik *et al.* (2017) (24) giving three pseudo biological replicates. RNA was extracted from each pseudo biological replicate and split into two and prepared as poly(A) RNA and Total RNA libraries. Raw sequencing reads were downloaded from the Gene Expression Omnibus (GEO) (25, 26) under accession number GSE64098. 12 libraries were examined for this dataset.

*Human immune cells* were sequenced by Linsley *et al.* (2014) (27) using a poly(A) RNA protocol; GEO accession number GSE60424. RNA samples were taken of whole blood and 6 immune cell subsets, including pure populations of neutrophils, monocytes, B cells, CD4+ T cells, CD8+ T cells and natural killer (NK) cells. 134 libraries were examined for this dataset.

*Mouse mammary cells* from female virgin mice with additional samples from mammosphere and the CommaD-*β*Geo (CommaD-bG) cell line were sequenced in a study by Sheridan *et al.* (2015) (28) to obtain poly(A) RNA libraries; GEO accession number GSE63310. Mammary cell populations include mammary stem cell-enriched basal cells, luminal progenitor-enriched (LP) and mature luminal-enriched (ML) cell populations. 19 libraries were examined for this dataset.

### Genomes and gene annotations

FASTQ files containing raw sequencing reads were aligned to the human *hg38* genome or mouse *mm10* genome using *subjunc* (29) with default parameters in the *Rsubread* software package (30). GENCODE’s main *Comprehensive gene annotation* file in GTF format was downloaded from https://www.gencodegenes.org for human (Release 27) and mouse (Release M12). GENCODE annotation was chosen over other annotations, such as RefSeq or Ensembl, because it is more inclusive and less stringent in terms of gene definitions and exon boundaries and also includes the annotation of long non-coding RNAs (8). Annotation files were simplified by taking the union of two or more overlapping exons from transcripts of the same gene. The adjustment provides a simplification of genomic positions on each strand, such that each position is classified as belonging to “exon”, “intron” or otherwise outside of an annotated gene. Three resultant annotation files were saved in standard annotation format (SAF) – exon annotation, intron annotation (region between exons), and genebody annotation (region spanning first to last exon). Scripts to process annotation files are available at https://github.com/charitylaw/Intron-reads, together with scripts for all other data analyses and supplementary figures.

### Intron and exon counts

Aligned reads were summarised by *featureCounts* (31) using exon annotation and genebody annotation separately to get gene-level *exon counts* and genelevel *genebody counts* respectively. In both counting strategies, reads that overlap features in multiple genes are not counted – exons in exon counting and genebody in genebody counting. For example, a read overlapping exons in two different genes is not counted towards exon counts, but a read overlapping an exon in one gene and an intron in another gene is counted towards exon counts; they do not contribute towards genebody counts in both cases since they overlap multiple genebodies. In this way, it is possible for exon counts be to greater than genebody counts for genes in some libraries. Gene-level *intron counts* are calculated by subtracting exon counts from genebody counts. For genes with exon counts that are greater in genebody counts, gene-level intron counts are adjusted to zero. Across our datasets, this occurred in ≈ 15% of genes where exon counts were greater than gene-body counts by a median value of −8. Intron counts represent the gain in information when summarising reads across the whole genebody relative to exon regions only. Whilst there are other count strategies, such as counting exon-intron boundary reads separately or towards intron counts, we take this approach since our interest is in assessing whether the intron reads that are not typically used contain additional signal.

### Coverage patterns

Coverage was estimated using our *superintronic* package for each BAM file for poly(A) RNA and Total RNA HCC827 human cell lines via the *Rsamtools* package (32, 33). Intron and exon regions were constructed for each gene from the GENCODE v27 annotation GTF. Genes in the annotation were restricted to protein coding genes on reference chromosomes. Genes that overlapped any other gene were also removed to reduce coverage ambiguity. Intron and exon regions from the set of genes were intersected with estimated coverage scores from BAM files for each library. This gives us an estimate of the number of bases covered at a given coverage score for each intron and exon region within a gene and sample. Coverage scores within each sample were transformed to log_2_-scale using an offset of 0.5. Genes were then further filtered if they were not expressed in the poly(A) RNA protocol (requiring at least a count of 3 in intron and exon regions). A total of 3,262 genes were examined and categorised as short, regular or long (roughly 1,087 genes in each category) by splitting the length of each gene into three bins by tertiles. Log-transformed coverage scores were then normalized by dividing by the maximum log-coverage value of each gene, giving *relative log-coverage* values. This stabilizes the variance of coverage values and quantifies the relative coverage change within genes.

To summarise coverage patterns across multiple genes, each gene was separated into 20 windows using the *GenomicRanges* package (33). We then found where windows intersected our relative log-coverage values using an overlap join from *plyranges* (34). For each sample and within each gene, we compute the *mean relative log-coverage* for each intron and exon region within a window based on position by taking the mean coverage value. We then compute across genes the mean of the *mean relative log-coverage* of each window for the short, regular, and long categorised genes. These were taken to represent general trends along the gene body (more specifically, the *mean of mean relative log-coverage* values).

### *IRFinder* method for differentially retained introns

*IRFinder* software version 1.2.4 (20) was applied to the comparison of HCC827 versus NCI-H1975 cell lines for poly(A) RNA libraries. We apply two separate models available – the Audic and Claverie (35) test for a small number of replicates and a generalised linear model (GLM) fit. In our analysis, the Audic and Claverie model interrogated 3,421 intron regions within 2,339 genes, and the GLM model interrogated 231,337 intron regions within 18,502 genes. Both models are fitted since the *IRFinder* manual available from https://github.com/williamritchie/IRFinder/wiki does not specify a specific test for our 3-versus-3 design, but suggests the Audic and Claverie test for 2-versus-2 design and GLMs for a 4-versus-4 comparison.

*IRFinder* detects IR for each library using the software’s in-built hg38 reference genome and relevant fastq files. In theory, the output here could be used to call retained introns within samples directly, however, there is no guideline on how to use the intron-summarised information. Instead results from each group are pooled to test for differences between cell lines using the Audic and Claverie method. The GLM method uses unpooled results to test whether the cell lines differ in their intronic-to-spliced reads ratio. *P*-values are output by both tests; adjusted *p*-values are calculated for GLMs only. In the Audic and Claverie analysis, only 2 genes met the expression criteria set by IRFinder; other genes were marked as having intron expression in one sample only.

### *IsoformSwitchAnalyzeR* method for differentially retained introns

The Bioconductor package *IsoformSwitchAnalyzeR* version 1.7.0 (23) was applied to human cell lines to detect differentially retained introns between the cell lines in poly(A) RNA libraries. The vignette of *IsoformSwitchAnalyzeR* suggests that best results are obtained when performing read quantification via pseudo alignment, so first we ran *kallisto* version 0.44.0 (5) in quantification and single-end mode on our libraries with the flags `-single -l 175 –s 20`. Quantified isoforms were imported using *IsoformSwitchAnalyzeR* and filtered out from downstream analysis if they did not meet the following criteria: they belonged to a protein coding gene, their gene expression was greater than 1 transcripts per million (TPM), the individual isoform expression was greater than 0.1 TPM. IR events were analysed with the `isoformSwitchTestDEXseq` function with a false discovery rate (FDR) cutoff of 5% and the default setting of isoform usage cutoff of 10%. After this step was completed we tested for IR events with `analyzeIntronRetention` and `analyzeSwitchConsequences` functions. This resulted in 502 genes detected with differential IR events at an FDR cutoff of 5% with between the poly(A) RNA and Total RNA libraries.

## Results

### Intron reads are prevalent across datasets

Taking a conservative approach, we quantify the number of intron reads that map entirely to an intron of a gene, excluding those that overlap an exon-intron boundary. Gene-level intron counts represent the extra counts one may obtain from within a gene when looking outside of annotated exons. The proportion of reads contributing to gene-level intron counts ranges from 2% to 14% with a mean value of 7% for poly(A) RNA libraries across three datasets examined (Figure 1a). A greater proportion of reads contribute towards gene-level exon counts, ranging from 57% to 78% with a mean of 69% (Figure 1b). Despite the relatively small proportion of intron reads, they amount to hundreds of thousands to millions of reads per library under typical sequencing protocols. For a library of size 30 million, the number of intron reads is approximately 2.1 million (using the mean value of 7%).

**Fig. 1.**
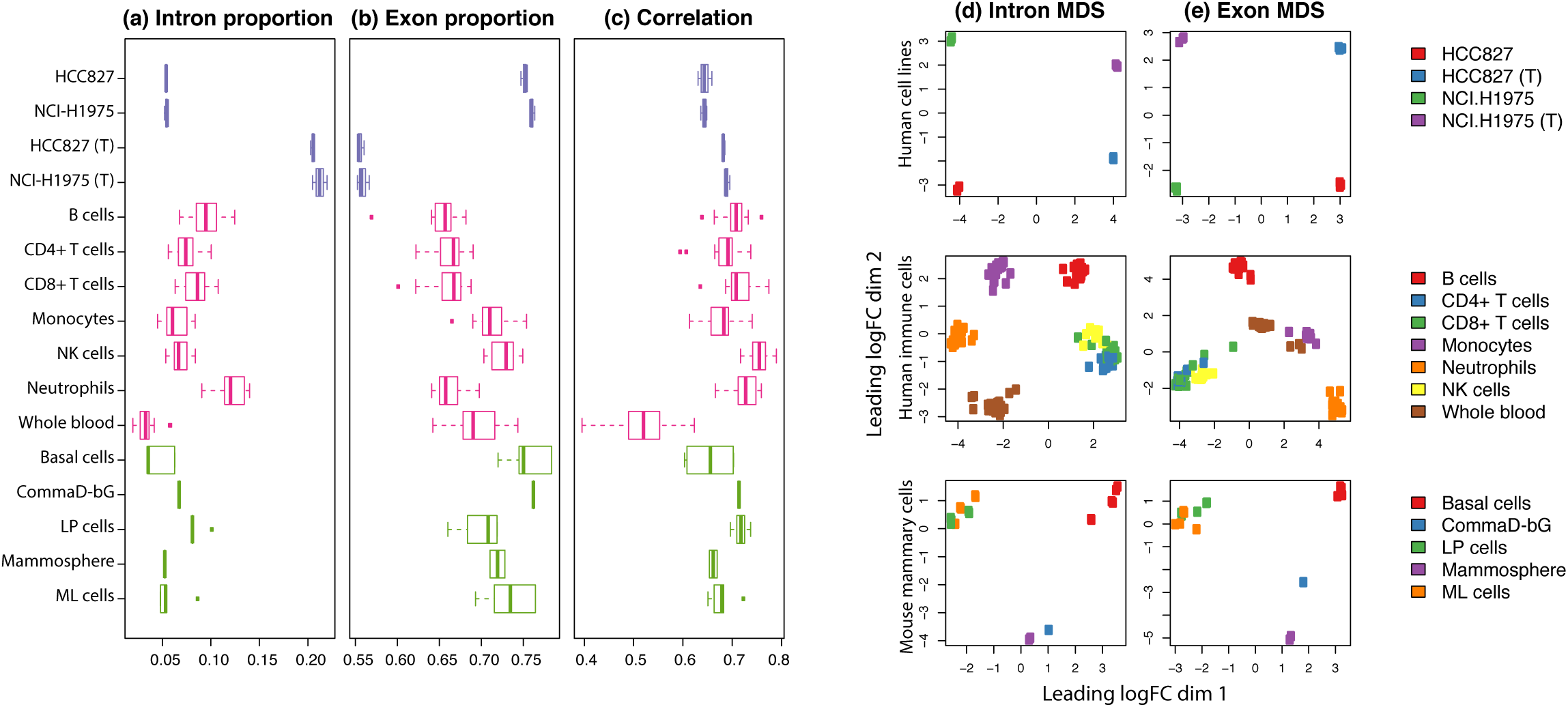
With libraries separated by biological and experimental groups, various statistics are summarised as boxplots across the datasets (distinguished by color – purple for human cell lines, pink for human immune cells, and green for mouse mammary cells) and Total RNA samples labelled with a ‘(T)’. (a) Proportion of reads assigned to intron and (b) exon counts. (c) Pearson correlation of gene-level exon log_2_-counts (log-counts) and gene-level intron log-counts. Log-counts are calculated for genes expressed (count of 3 or more) in both intron and exon regions, using an offset of 1. (d) MDS plots of log_2_-counts-per-million (log-CPM) values calculated using an offset of 2 for gene-level intron counts and (e) gene-level exon counts for each of the three datasets. MDS plots were created using *limma*’s (36) `plotMDS` function based on the top 500 most variable genes.

A higher proportion of intron reads are found in Total RNA libraries in comparison to poly(A) RNA libraries, as noted in prior studies (11, 14). The mean proportion of reads contributing to intron counts and exon counts for Total RNA libraries in human cell lines is 21% and 56%, respectively – a profound difference of roughly 15% more intron counts and 20% fewer exon counts when compared to corresponding poly(A) RNA samples. This equates to approximately 6.3 million intron reads for a library of size 30 million. Intron and exon read proportions are fairly consistent for libraries within the same biological and experimental groups (such as within cell lines, cell types, tissues, RNA library protocol). Some variation in read proportions can be observed for different biological (e.g. neutrophils versus whole blood) and experimental (e.g. poly(A) RNA versus Total RNA HCC827 cell line) groups. Within libraries, exon log-counts are positively correlated with intron log-counts (Figure 1c).

### Intron reads are informative and contain biological signal

In DGE analyses, plots of principle components analysis and multi-dimensional scaling (MDS) methods are commonly created from exon counts to provide an overview of the similarities and differences in transcriptional profiles in an unsupervised manner. To determine whether intron reads contain any biological signal, we applied MDS methods to intron counts instead. Samples cluster by experimental and biological groups in intron MDS plots across all datasets (Figure 1d) indicating that intron reads are informative, rather than a result of sequencing noise. As expected, samples also cluster by experimental and biological groups in exon MDS plots (Figure 1e).

Extraordinarily, the scales observed in the first and second dimensions of intron MDS plots are comparable to that of exon MDS plots even though there are roughly ten times fewer intron counts than exon counts. The distance between points on each plot give an indication of the typical log_2_-fold change (logFC) between samples in the top 500 most variable genes in each set of counts. In other words, the typical logFC between samples are similar for intron and exon counts.

Note that the first dimension of separation accounts for a larger proportion of variation in the data than the second dimension. The MDS plots for human cell lines indicates that intron read signal is significantly influenced by RNA selection protocols. The first dimension in the intron plot separates samples based on RNA selection method and accounts for 48% of the variation in the intron counts, whilst the second dimension relates to to cell line identity and accounts for 18% of variation in the data. This is in contrast to the exon plot where the RNA selection protocol (second dimension) accounts for 26% of the variation in counts, and cell lines identify (first dimension) accounts for 37%. The type of RNA selection protocol used in library preparation has a greater influence on intron reads than exon reads.

### Characteristics of counts

The human cell line dataset allows us to explore count differences between library preparation methods. Gene-level exon log-CPM values are similar between poly(A) RNA and Total RNA libraries and have a very strong positive correlation (Figure 2a). Gene-level intron log-CPM values are also positively correlated but counts tend to be greater in Total RNA than poly(A) RNA libraries (Figure 2b). Log-CPM values were calculated using an offset of 2 and by setting the library size as the sum of counts from exons and introns (the same is done for log-RPKM values). This allows adjustment of intron and exon counts by the same sequencing depth per library, rather than an intron- or exon-specific proportion of the original sequencing depth. Within libraries, we determined whether the same genes expressed intron and exon signal (had reads overlapping both intron and exon regions), or whether intron and exon signal came from different genes. We found that 56% of expressed, multi-exonic genes were expressed in both intron and exon regions on average across human cell line libraries. Expressed genes have a count of 3 or more in exon and/or intron counts, and genes expressed in both intron and exon regions have a count of 3 or more in both intron and exon counts. This means that the majority of expressed, multi-exonic genes contain both intron and exon signal simultaneously. 32% of expressed, multi-exonic genes were expressed in exon regions only (exon count ≥ 3, intron count ≤2), and 13% of genes were expressed in intron regions only (exon count ≤2, intron count ≥3) on average.

**Fig. 2.**
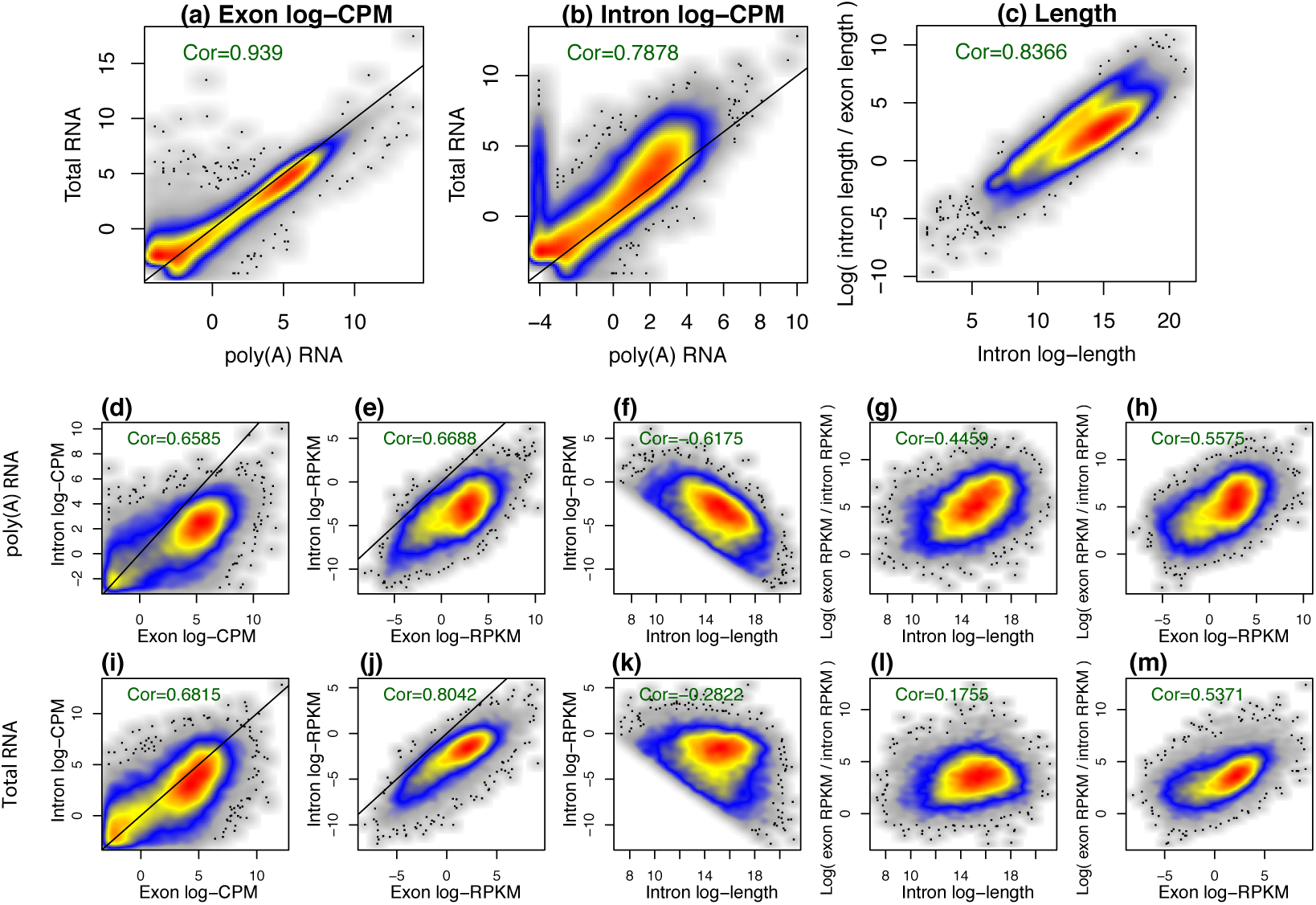
For HCC827 human cell line R1 sample (a) RNA protocols are compared for gene-level exon log-CPM and (b) gene-level intron log-CPM values. Genes that are expressed (log-CPM *> -*3, which roughly translates to 3 or more counts in each library) in either of the libraries under comparison are included in the plots. Across all plots, each point represents a gene; density of points are distinguished by varying color shades, increasing from grey, blue, yellow, through to red; extreme points are marked by black dots; a line of equality is drawn within comparisons where positive correlation is expected; and Pearson correlation values are marked in green within each plot. (c) On a log_2_-scale, the relative length of introns compared to exons are plotted against the length of introns. The exon length of a gene is calculated as the sum of all exon regions as defined in the exon annotation; similarly for intron length. Log-values are calculated using an offset of 0.001. Within the HCC827 R1 poly(A) RNA library, intron and exon (d) log-CPM and (e) log_2_-reads per kilobase million (log-RPKM) values are compared to each other, using an offset of 2. (f) Coverage (in log-RPKM) is compared to length for intron regions; and (g) relative coverage between exon and intron regions is compared to length of intron regions, and (h) to exon coverage. In panels (d) to (h), only genes that are expressed in both intron and exon regions (log-CPM *> -*3) are included; similarly for the equivalent HCC827 R1 Total RNA plots in panels (i) to (m). The plots shown here are for HCC827 R1 libraries, however, similar plots were observed for all HCC827 and NCI-H1975 cell lines samples (Supplementary Materials).

To understand the nature of intron counts in relation to exon counts, we focus on the set of genes that are expressed in both regions, noting that genes with large total intron length (sum of all bases annotated as not belonging to an exon) have proportionally much longer intron regions than exon regions (Figure 2c). Within poly(A) RNA libraries, intron and exon log-CPM and log-RPKM values are positively correlated (Figure 2d and e), where log-CPMs provide a reflection of the size of counts used as inputs to many analysis methods and log-RPKMs are adjusted for length differences between intron and exons regions within the same gene. Figure 2e shows that intron coverage tends to be lower than exon coverage within the same gene, but the relative difference is quite stable across the genes. The median difference between gene-wise exon log-RPKM and intron log-RPKM values is ≈ 5.1 across all poly(A) RNA HCC827 and NCI-H1975 cell line libraries, such that gene-wise exon coverage is roughly 34 times greater than intron coverage on average. Average intron coverage is affected by total length of intron regions in genes, such that genes with longer intron regions tend to have lower log-RPKM values (Figure 2f). Also, relative coverage of exons over introns increases for genes as total length of intron regions increases (Figure 2h), and as exon log-RPKM increases (Figure 2h).

Similar trends are observed in Total RNA samples (Figure 2i-m), though it is worth noting that intron and exon counts are similar in count size and dynamic range for Total RNA libraries, and have stronger correlation of log-CPM and log-RPKM values between introns and exons. For Total RNA libraries, exon coverage is roughly 10 times greater than intron coverage on average (median difference between gene-wise exon log-RPKM and intron log-RPKM values is approximately 3.3 across all Total RNA HCC827 and NCI-H1975 cell line libraries).

### Intron reads are predominantly from pre-mRNA

The notion that intron reads originate from pre-mRNA molecules rather than genes with retained introns is supported by the observation that gene-level exon counts tend to have a strong positive correlation with intron counts across all genes (Figure 1c and Figure 2d and i), and that that a large proportion of expressed, multi-exonic genes express intron and exon signal simultaneously. Assuming that IR is generally not widespread across all genes, a weak positive correlation is expected between intron and exon counts if intron reads were predominantly coming from genes with retained introns.

Coverage patterns across the genebody of multi-exonic genes provide further evidence that intron signal is predominantly from pre-mRNAs with unspliced or partially spliced introns, where intron reads tend to be uniformly distributed in Total RNA libraries and increasing gradually towards the 3’ end of genes in poly(A) RNA libraries (Figure 3a). Total gene length appears to play a part in intron coverage patterns. Genes under examination were catergorised by total gene length (length from first base in first exon to last base in last exon) such that a third were considered to be *short, regular* and *long* genes each. Coverage patterns were similar between poly(A) RNA and Total RNA libraries for short genes, whilst patterns differed substantially for regular and long genes. Exon coverage patterns were similar for poly(A) RNA and Total RNA libraries (Figure 3b).

**Fig. 3.**
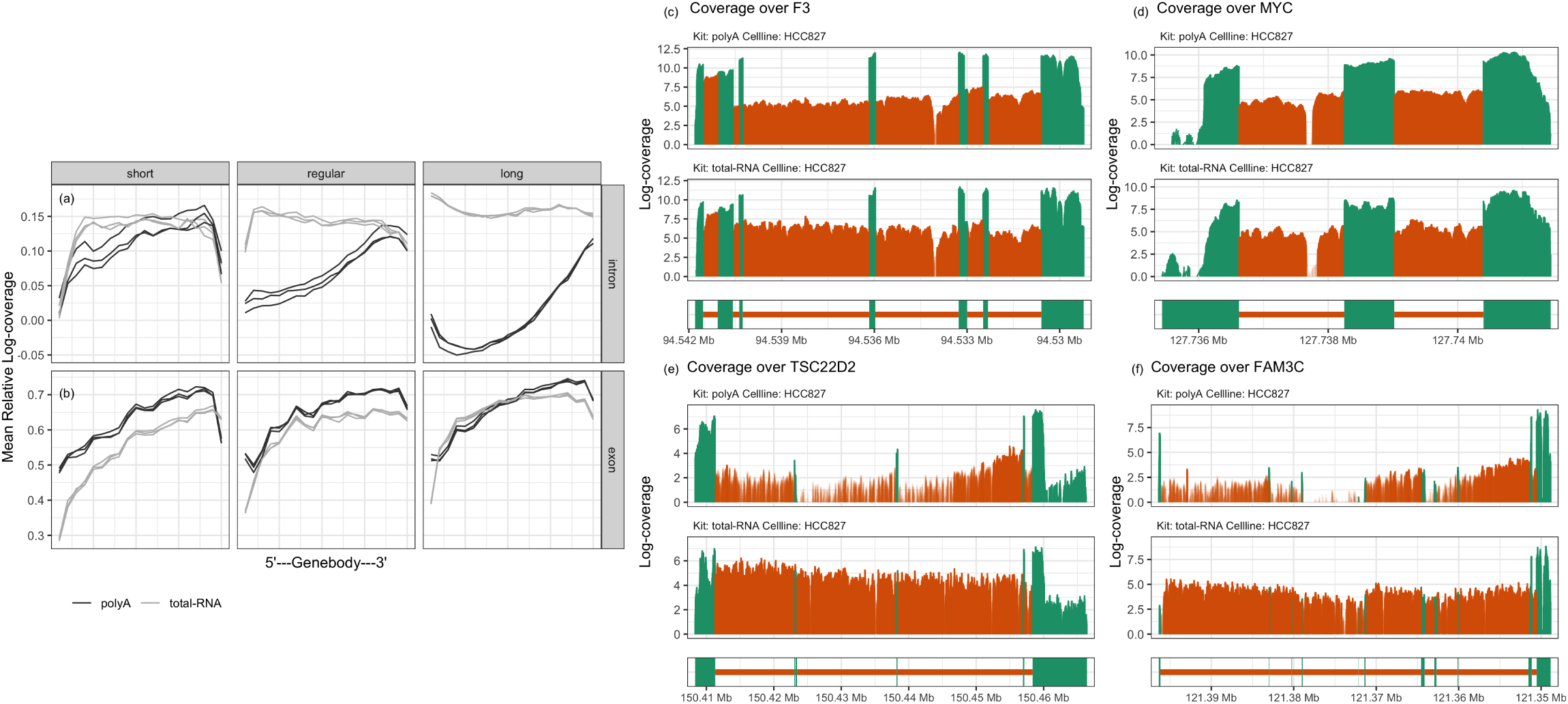
Coverage of intron and exon regions across genebody. (a) Coverage patterns of intron regions across body of genes, separated into groups based on gene length. Coverage is represented by the mean of mean relative log-coverage, where relative log-coverage is calculated as local log-coverage divided by maximum log-coverage in a given gene. HCC827 cell line samples are depicted with black lines representing poly(A) RNA libraries R1, R2 and R3, and grey lines representing the corresponding Total RNA libraries. (b) Coverage patterns of exon regions across body of genes. (c) to (f) Log-coverage in exon regions (green) and intron regions (orange) in poly(A) RNA R1 library (top) and Total RNA R1 library (bottom) are displayed for two short genes, (c) MYC and (d) F3, and two long genes, (e) FAM3C and (f) TSC22D2. These genes were selected based on having a high median intron log-coverage. Intron regions with high expression (log-coverage greater than 3) are highlighted by a shade of darker orange.

Coverage patterns are represented by *mean of mean relative log-coverage* values. Briefly, the mean coverage (or *mean relative log-coverage*) is calculated for 20 sections along each gene and the mean of the each of the sections is calculated across multiple genes giving the *mean of mean relative log-coverage*. Since we are interested in relative changes in coverage within a gene, *relative log-coverage* is examined rather than absolute values. *Relative log-coverage* refers to coverage values that have been log-transformed and divided by the maximum log-value. Further details can be found in the Materials and Methods section.

Confirming the same patterns for individual genes, coverage profiles of two short genes MYC and F3 are observed to be similar between RNA protocols (Figure 3c and d). In contrast, the coverage profiles of two long genes FAM3C and TSC22D2 differ at the 5’ end where poly(A) RNA libraries are observed to have deflated intron coverage relative to the 3’ end, as well as relative to Total RNA libraries (Figure 3e and f). Reduction in read coverage at 5’ introns for poly(A) RNA libraries explain why poly(A) libraries are observed to have relatively low proportions of intron reads (Figure 1a) and dynamic range in log-CPM values relative to Total RNA libraries (Figure 2d and i).

Relative gene-level contribution by pre-mRNAs is low compared to mature mRNAs for the majority of genes, as reflected in low intron log-RPKM relative to exon log-RPKM values (Figure 2e and j). If intron and exon log-RPKM values provide an estimate of the relative proportions of pre-mRNA and mRNA molecules captured, then on average roughly 1 pre-mRNA molecule is captured for every 10 molecules in a sequencing experiment (since exon coverage is roughly 10 times greater than intron coverage in Total RNA libraries, and Total RNA libraries have uniform intron coverage). Unless the sequencing experiment is carried at very high depths, intron signal may not be detected for genes with relatively short intron regions since pre-mRNA levels are low, whilst genes with long intron regions have greater ability to accumulate sequencing reads over the gene.

### DGE analyses of transcriptional activity using intron and exon counts

Classical DGE analyses are performed on gene-level exon counts where associated reads originate from mRNA and pre-mRNA molecules. We propose a method which also includes intron counts into the analysis to measure changes in early transcriptional activity. We call our method *index*, **in**tron **d**ifferences to **ex**on, a DGE method categorising genes by significance and directional changes in intron and exon counts. Relative to a classical DGE analysis which requires gene-level exon counts and some information about the experimental design and comparisons of interest, *index* simply requires an addition of gene-level intron counts. *Index* is an R package which is available to download and install at https://github.com/Shians/index.

An *index* analysis is carried out on genes that are expressed in both intron and exon regions – sufficiently large intron and exon counts are selected by the *filterByExpr* function in *edgeR* (37, 38). Trimmed mean of M-values (TMM) normalisation (39) is then carried out on intron and exon count sets separately using the combined library size, sum of both prefiltered intron and exon counts together, for samples. This is a variation on the standard library size calculation which sums counts within a single count set. Intron and exon counts evaluated based on single library sizes will be affected by intron and exon read proportions (Figure 1a and b) which vary between samples, groups and experiments, and gives a poor estimate of original sequencing depth. The combined library size is used also for downstream calculations, such as in obtaining log-CPM values by *voom* (40).

DE genes are obtained for intron and exon counts separately following a standard *limma-voom* pipeline (41) where log-CPM values and variables of interest are modelled on a Normal distribution with precision weights calculated by *voom*. Moderated *t*-statistics are calculated for each gene using Empirical Bayes’ methods (36, 42) and *p*-values are adjusted for multiple testing by controlling the FDR (43). Genes with an adjusted *p*-value of less than a nominal cutoff are considered to be DE.

*Index* categories are formed based on significance in intron and exon counts: **+** for genes up-regulated in both intron and exon counts, **-**for down-regulated in intron and exon counts, **exon+** and **exon**-for up- and down-regulated in exon counts only, **intron+** and **intron-**for up- and down-regulated in intron counts only, **mixed+-**for up-regulated in exons and down-regulated in intron counts and **mixed-+** when in the opposite direction, and **0** for no significant difference in either exon or intron counts.

The *index* software performs analysis on intron and exon DGEList objects (a native object of *edgeR*) to classify genes into the respective *index* categories. *Index* outputs the categories assigned to each gene, the tables containing *limma* differential expression analysis for introns and exons, and other data that is used to create plots from the software’s plotting functions. This allows the *index* analysis to be easily performed on any dataset where intron and exon counts can be separately obtained.

### *Index* analysis of human cell lines and immune cells

An *index* analysis comparing NCI-H1975 versus HCC827 cell lines in Total RNA libraries reveals that the majority of genes are DE in the same direction between intron and exon counts – 2,406 genes up-regulated (+) in NCI-H1975 and 2,464 down-regulated (-) using an adjusted p-value cutoff of 0.01 (Figure 4a-c). Genes DE by exon counts only form the second biggest group, with 989 genes up-regulated (exon+) and 914 genes down-regulated (exon-) in exon counts. Interestingly, these genes tend to have short intron regions (Figure 4d). There are 547 genes up-regulated (intron+) and 459 genes down-regulated for intron counts only (intron-), where genes tend to have relatively long intron regions. Similarly, genes DE in opposite directions also have relatively long intron regions – a small group of 25 genes up-regulated in exon counts but down-regulated in intron counts, and 29 genes up-regulated in intron counts but down-regulated in exon counts. The analysis was carried out on 11,608 genes after lowly expressed genes were filtered out.

**Fig. 4.**
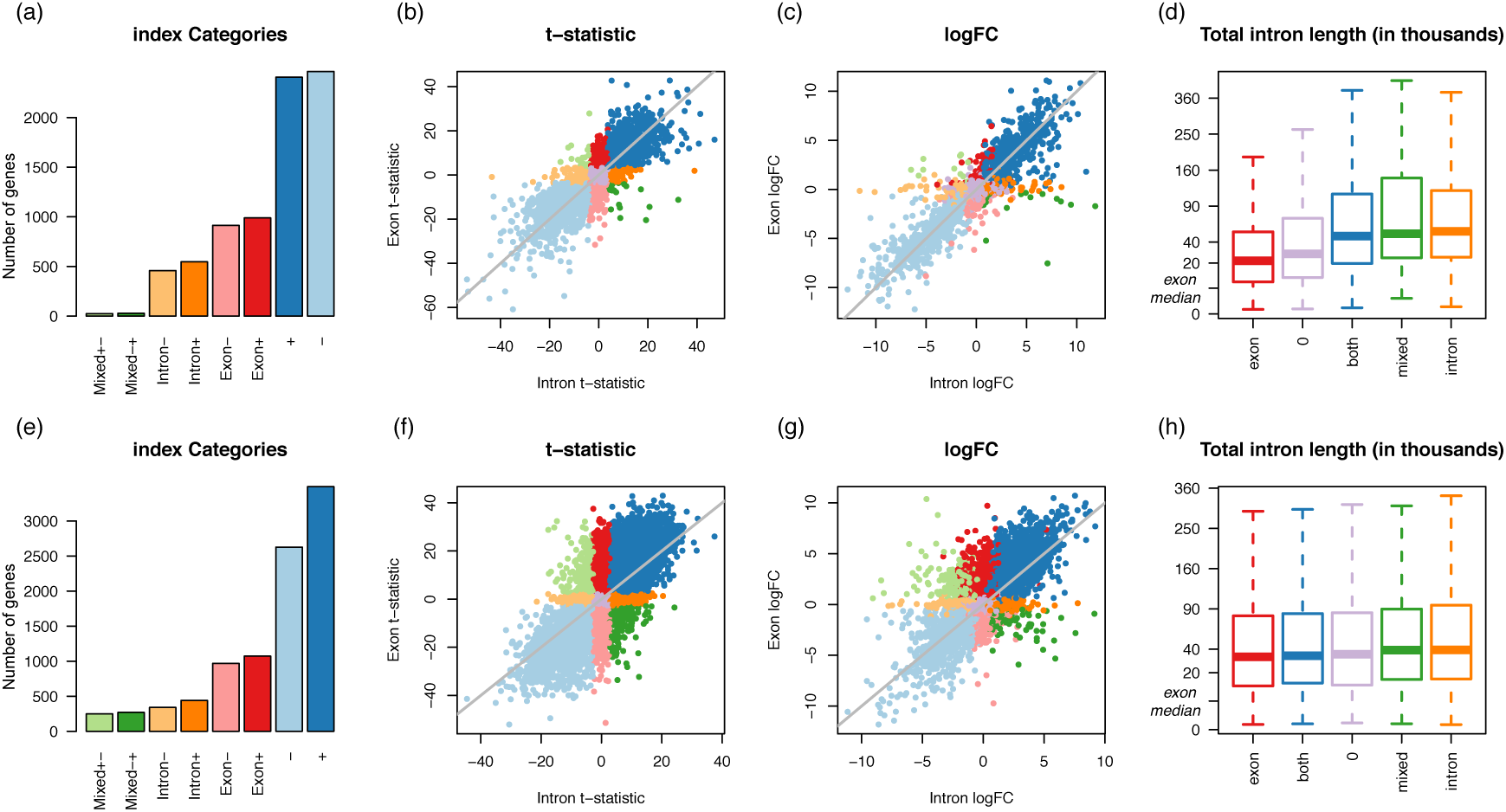
*Index* DGE analysis of intron and exon counts. (a) Number of significant genes in *index* categories for NCI-H1975 versus HCC827 cell lines in Total RNA libraries, with (b) t-statistics from exon counts plotted against those from intron counts and (c) logFC values from exon counts plotted against those from intron counts, where colors are associated with different *index* categories. (d) Distribution of total intron length where *index* categories are combined, such that genes that are up-regulated or down-regulated for both intron and exon counts are reassigned to *both*; *exon* for genes up- or down-regulated for exon counts only, similarly for *intron* and *mixed*. A square-root scale is used along the vertical axis, with the median total exon length (5,126 bases) marked as a reference. (e) to (h) Similar plots are displayed for a comparison of monocytes and neutrophils in the immune cells dataset.

Identical analysis comparing monocytes versus neutrophils in the immune cells dataset similarly reveals that the majority of genes are DE in the same direction between intron and exon counts – 3,491 genes are up-regulated (+) in neutrophils relative to monocytes and 2,628 down-regulated (-) using an adjusted p-value cutoff of 0.01 (Figure 4e-g). Again, genes DE by exon counts only form the second biggest group, with 1073 genes up-regulated (exon+) and 969 genes downregulated (exon-) for neutrophils in exon counts. These genes again tend to have shorter intron regions, but the difference in length between *index* catergories is subtle compared to that of cell lines (Figure 4h). There are 441 genes up-regulated (intron+) and 340 genes down-regulated (intron-) for intron counts only, where genes again tend to have longer intron regions. The number of genes DE in opposite directions between intron and exon counts is larger for this comparison than in cell lines, with 249 genes up-regulated in exon counts but down-regulated in intron counts (mixed+-), and 269 genes down-regulated in exon counts but up-regulated in intron counts (mixed-+). The analysis was performed on 10,789 genes after lowly expressed genes were filtered out

### Index analysis detects additional DE genes

The *index* DGE analyses demonstrate that transcriptional changes detected by exon counts are similar to those detected by intron counts. This is expected since exon counts represent mRNA and pre-mRNA levels, whilst intron counts represent pre-mRNA levels. For most genes, similarity between intron and exon logFCs (Figure 4c and g) indicate that pre-mRNA and mRNA levels are simultaneously up- or down-regulated at similar proportions between groups.

Assignment of genes into different *index* categories is associated with total intron length of a gene (Figure 4d and h), such that genes DE for exon counts only tend to have relatively short intron regions. Naturally, these genes are unlikely to accumulate high intron counts due to low coverage and short region lengths, thus lacking power during statistical testing. On the other hand, genes DE for intron counts only tend to have relatively long intron regions; due to their length they are able to accumulate high intron counts even if coverage levels are low, giving them a power advantage when testing for differential expression.

In other words, exon+ and exon-genes may also contain changes in intron regions even though they remain undetected. Alternative explanations for observing significant changes in exon counts only are less likely. For example, that there no pre-mRNAs observed, or that pre-mRNA levels are consistent between groups. The former is contradicted by Figure 2f and k which shows high intron coverage for genes with short intron regions, and the latter is unlikely to be trait specific to genes with short intron regions. Similarly, intron+ and intron-genes may also contain changes in exon regions even though they remain undetected.

If intron counts represent pre-mRNA levels, then any change observed between groups in intron counts should also be reflected in exon counts. However, exons are unlikely to accumulate high counts over its relatively short regions if pre-mRNA (and mRNA) levels are very low. If genes have retained introns or differentially retained introns in one group versus another, it is also possible for genes to be detected as DE in intron counts. Intron+ and intron-genes can be compared against a list of genes detected with retained introns (see *superintronic* in next section). Given that significance is influenced by total intron length, it is possible that exon+, exon-, intron+ and intron-genes may be reclassified into + and - *index* categories if sequencing was performed at greater depths.

Mixed+- and mixed-+ genes form a relatively small set of genes relative to other *index* catergories. Biologically, a simultaneous increase in pre-mRNA levels and decrease in mRNA levels between two groups can induce changes in opposing directions between intron and exon counts. The results suggest that this occurs relatively infrequently, however, when it occurs *index* is able to differentiate these genes from those where the direction of change is consistent between pre-mRNA and mRNA molecules.

An *index* DGE analysis adds an extra layer of information by overlapping intron and exon results, where additional DE genes are detected that are not observed in a classic DGE analysis alone. The *index* method has increased power in genes with long intron regions, where high counts can be detected for low coverage genes. A classic DGE analysis, by *limma-voom* or like methods, followed by an *index* DGE analysis allows researchers to make use of a larger proportion of reads that are already sequenced and available to them to detect additional DE genes.

### *Superintronic*: a novel method for detecting genes with IR

Considering IR also as another possible source of intron reads we propose a new method using our *superintronic* software to find retained introns, with the assumption that most intron reads do not point to IR but pre-mRNA instead. *Superintronic* is an R package that is available to download and install at https://github.com/sa-lee/superintronic. It extends the *plyranges* Bioconductor package (34) for genomics data analysis by estimating and aggregating coverage scores directly from BAM files. *Superintronic* requires BAM files, gene annotation and information on experimental design as input.

Our software records the per base coverage over intron and exon regions of each gene, with the option of storing these per sample or summarised over variables in the experimental design such as by biological group or by RNA protocol. Coverage scores are normalised using a log_2_-transformation with an offset of 0.5 to get log-coverage values for which intron and exon summary statistics are constructed for each gene (described below). Within *superintronic*, a suite of visualization tools to construct coverage plots for genes with intron and exon structures and scatter plots are provided.

### *Superintronic* finds genes with IR-like coverage profiles in human cell lines

Using *superintronic*, poly(A) RNA HCC827 cell lines were examined for genes with IR after selecting genes in the hg38 reference that were protein coding, did not overlap any other gene and were placed on the main contigs – a total of 6606 genes. These genes were then split into intron and exon regions and intersected with the coverage of each sample. Per gene intron and exon summary statistics, mean and standard deviation, were computed on log-coverage values. Genes enriched for IR-like coverage profiles were selected based on a set of thresholds – genes had an average exon log-coverage of greater than 2 (corresponding to the mean of average exon log-coverage values across all genes), a standard deviation of intron log-coverage greater than 1.5 (corresponding to the mean of exon standard deviation values across all genes), and genes with a large number of intron bases with log-coverage greater than 2 (top 1% of genes). The thresholds were chosen after examining distributions of the summary statistics (Supplementary Materials). These filters select for “expressed” genes, where for a substantial number of intron bases its coverage is much higher than other intron features within the same gene while having similar expression levels to the exon features. Forty-three genes met these criteria (Supplementary Materials), where a manual check of coverage profiles revealed that 36 genes indeed appear IR-like. We highlight 3 of these genes in Figure 5. The coverage of remaining 7 genes appear to be more pre-mRNA-like, with large variation in intron coverage, where at its peak it is expressed similarly to exon features. We use poly(A) RNA samples in our analysis for consistency with previous studies on IR. In theory, Total RNA samples may be more appropriate for this exercise since it is less biased towards the 3’ end, where high 3’ intron coverage as a result of 3’ bias in poly(A) RNA libraries can be mistaken as a 3’ retained intron. For this reason, we review both the coverage of poly(A) RNA and Total RNA samples in our supplementary materials for all selected genes to ensure that retained introns found in poly(A) RNA samples are not an artifact of RNA protocol.

**Fig. 5.**
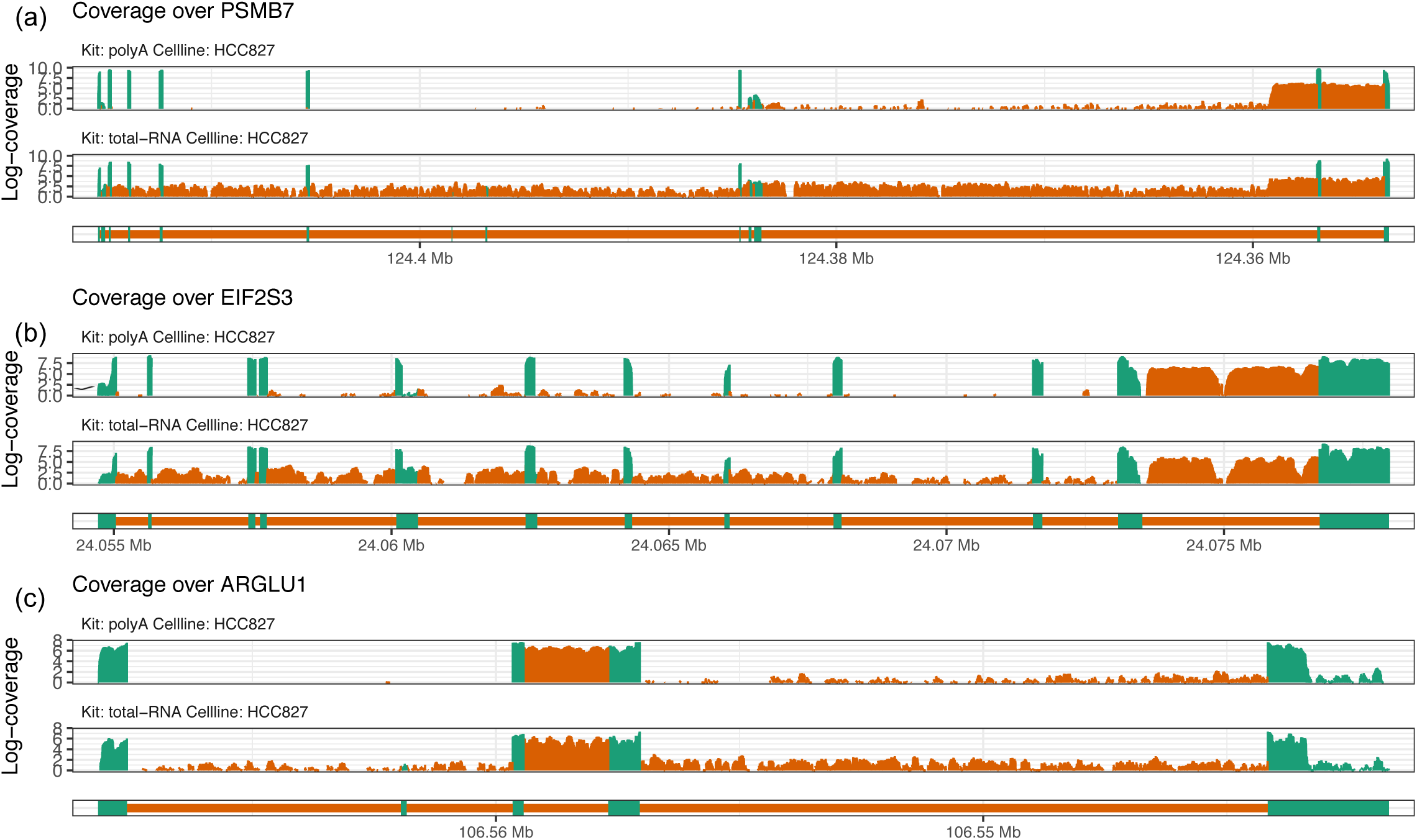
*Superintronic* selects genes with IR-like coverage profiles in poly(A) RNA HCC827 cell lines. Genes (a) PSMB7, (b) EIF2S3, and (c) ARGLU1 are highlighted out of 43 genes selected. Whilst the analysis was carried out on poly(A) RNA libraries, coverage is shown for both poly(A) RNA HCC287 and Total RNA HCC287 samples to ensure that results are not an artifact of RNA protocol. Coverage is oriented from 5’ to 3’, with exon regions colored green and intron regions colored orange.

Our method differs from existing methods such as *IRFinder* and *IsoformSwitchAnalyzeR* in that it detects retained introns in individual conditions, rather than differences between two conditions. Naturally, the detection of differentially retained introns within a gene should firstly include a retained intron in at least one condition, and secondly contain differences in the expression of the intron. Since detection either of these steps are non-trivial, we believe that it is more important to focus on detecting retained introns correctly. IntEREst (22) is another method that detects IR in genes. We did not test this method as the authors have shown that results from IntER-Est overlap considerably with that of *IRFinder*. The Whippet software (21) has a module for identifying transcripts with retained introns, however, we were unable to install the software with several attempts.

If differentially retained introns are of interest, one could simply run *superintronic* on two conditions separately and compare lists of IR-like genes between groups. We found that differences between the lists of IR-like genes were concordant with *index* results for genes DE by intron counts, as expected (Supplementary Materials). For example, 14 genes were uniquely selected as IR-like in Total RNA HCC827 cell line when compared to Total RNA NCI-H1975 cell line using *superintronic*, where 13 of those genes were also found to be DE in intron counts by *index*, and in the expected direction. Similarly, 19 genes were uniquely detected as IR-like in Total RNA NCI-H1975 by *superintronic*, 16 of which were also detected as DE in intron counts by *index*. Additionally, 12 genes which were found to be IR-like in both cell lines appear to have varying intron expression according to *index* since it is found to be DE in intron counts.

### *IRFinder* and *IsoformSwitchAnalyzeR* detects pre-mR-NA-like genes

Although *superintronic* is not directly comparable to existing methods for detecting differentialy retained introns, we assess the performance of *IRFinder* and *IsoformSwitchAnalyzeR* on the human cell line data to see how they compare to *superintronic*. We make a comparison between poly(A) RNA HCC827 cell lines versus poly(A) RNA NCI-H1975 cell lines using both methods, with the assumption that genes detected to contain differentially retained introns should have at least a retained intron in one of the cell lines.

*IRFinder*’s Audic and Claverie small samples test detected differential IR in gene NBEAL2 (p-value of 0.0044) and CYR61 (p-value of 0.067), however their coverage profiles appear to be pre-mRNA-like and we could not identify the location of a retained intron in either conditions (Figure 6a). Other regions were deemed insignificant.

**Fig. 6.**
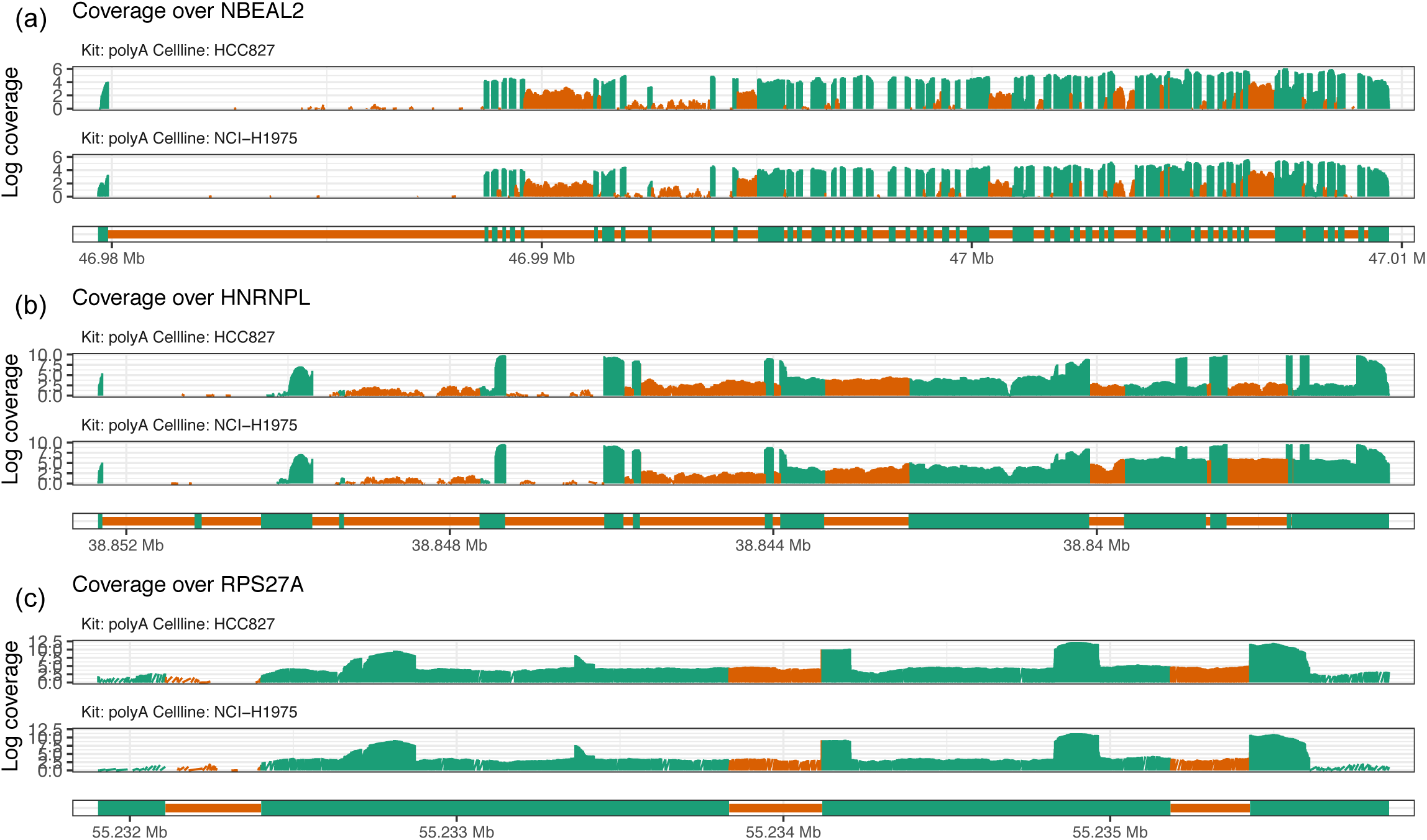
Genes with differentially retained introns between poly(A) RNA humam cell lines as detected by *IRFinder* and *IsoformSwitchAnalyzeR*, visualized as coverage plots facetted by cellline. (a) HNRNPL detected by *IRFinder* GLM test (b) NBEAL2 detected by *IRFinder* Audic and Claverie test and (c) HLA-B detected by *IsoformSwitchAnalyzeR* DEXSeq test. Coverage is oriented from 5’ to 3’, with exon regions colored green and intron regions colored orange.

An alternative method by *IRFinder* using GLMs detects 572 intron regions within 304 genes to contain differentially retained introns (5% adjusted p-value cutoff). Results between the two *IRFinder* methods are inconsistent, where genes in the more conservative method are not detected as significant under the GLM method (NBEAL2 and CYR61 have adjusted p-values of 0.84 and 1, respectively).

Under the GLM model, the 10 most significant regions as ranked by adjusted p-value and flagged as “clean” (excludes regions flagged by warning messages) are found across 6 genes with adjusted p-values ranging from 2.5 × 10^−17^ to 4.1 × 10^−9^. One of the top genes include HNRNPL (Figure 6b). As it was with the Audic and Claverie test, the GLM method detected genes that do not resemble the presence of IR in either of the cell lines, but instead appear pre-mRNA-like. Both methods appear to select for genes that have variable intron coverage, rather than retained introns

Likewise, we performed the same analysis comparing the poly(A) RNA HCC827 cell lines against poly(A) RNA NCI-H1975 cell lines for retained introns using *IsoformSwitchAnalyzeR*. Using an FDR of 5% we obtained 502 genes that were declared as having differentially retained introns, in which 255 isoforms were declared as a gain of IR, 470 isoforms were declared as a loss of IR and 24 isoforms were declared as a switch of IR. Upon inspection of coverage plots of the top ranked genes (designated a q-value < 0.01), we found many results to look pre-mRNA-like, similar to that of *IRFinder*. One of the top genes include RPS27A (Figure 6c). Coverage plots for other top genes detected by *IRFinder* and *IsoformSwitchAnalyzeR* can be found in Supplementary Materials.

Between the methods, 14 genes overlapped between results from *IRFinder*’s GLM method and *superintronic* run on both poly(A) RNA cell lines. *IsoformSwitchAnalyzer* overlapped with *IRFinder*’s GLM method by 15 genes, but there were no overlaps with *superintronic*.

## Discussion

The work presented in this paper provides a broad view of the characteristics and expression patterns associated with intron reads in model organisms, namely human and mice. Less studied organisms with poor and incomplete gene annotations may contain higher proportions of intron reads as a result of unannotated exons. In such cases, “intron reads” that are split reads can be useful for identifying new exons. *Index* and *superintronic* results can also be examined against a list of genes containing intron split reads to assess whether significant results are affected by unannotated exons. Additionally, any transcriptional changes in unannotated exons of annotated genes would be reflected in intron counts and could be detected in an *index* DGE analysis, and otherwise would be missed in a classic DGE analysis.

Using gold standard differential expression methods, *index* selects and categorises genes of interest based on p-values from moderated *t*-statistics that are adjusted for multiple testing and the direction of change. This is statistically more sophisticated than the *EISA* method which uses intron and exon logFCs alone and provides no prioritisation of genes of interest – this is, however, sufficient for their purpose of catergorising genes as sets, and determining whether the biological system under study is driven transcriptionally or post-transcriptionally as a whole. In contrast, *index* looks for DE genes using evidence from intron and exon counts. *Index*’s logFC plot (Figure 4c and g) is analogous to *EISA*’s main result and logFC plot.

The presence of pre-mRNA in poly(A) RNA libraries may be somewhat surprising since the RNA protocol is optimised for mRNA selection. However, pre-mRNA can be captured by both poly(A) RNA and Total RNA protocols since transcription from DNA to a primary RNA transcript, 3’ cleavage of the RNA molecule and polyadenylation can be completed before splicing is complete at the 3’ end because the splicing mechanism requires a relatively long processing time (44) – this is regardless of whether genes are co-(11, 45, 46) or post-transcriptionally spliced. Evidence supporting this includes the presence of poly(A)-positive molecules in the nucleus that are larger than final mRNAs in the cytoplasm (44). Handling of RNA in preparation for sequencing results in a degree of fragmentation of the original molecule regardless of the level of care taken during this process. This has minimal downstream effects on Total RNA libraries since 3’ and 5’ fragments are selected randomly. However, the selection of poly(A)-positive RNA molecules in poly(A) RNA libraries bias fragments at the 3’ end whilst 5’ fragments are lost in the process. This results in 3’ coverage bias in poly(A) RNA libraries (Figure 3) – demonstrated also by Lahens *et al.* (2014) (47) in their study on technical biases introduced during generation of sequencing libraries. Shorter genes with fewer and/or shorter introns are less affected by fragmentation than genes with long RNA molecules, thus coverage profiles are more similar between Total RNA and poly(A) RNA libraries for these genes (Figure 3). Read coverage in Total RNA libraries may provide a more accurate representation of the original RNA molecule than in poly(A) RNA libraries, especially in long genes. In poly(A) RNA libraries, the inflated 3’ exon expression and 3’ “background” intron signal may negatively impact on methods for *de novo* transcriptome assembly and transcript quantification. Total RNA libraries which have uniform “background” intron signal is easier to model in theory and should be better suited to such applications. Moreover, existing IR detection methods applied to poly(A) RNA sequencing libraries will have increased difficulty in interrogating introns residing towards the 5’ end of genes. (For the same reason, we also observe significant results tending towards towards the 3’ end of genes for both *IRFinder* and *IsoformSwitchAnalyzeR*.)

Compared to bulk RNA-seq, single-cell RNA-seq (scRNA-seq) data have much smaller library sizes and relatively high proportions of intron reads leading to much interest in the incorporation of intron reads in scRNA-seq data analyses (48, 49). Though yet to be tested, DGE analysis by *index* should theoretically perform well on scRNA-seq data since it increases the number of testable genes and libraries by increasing the amount of information used.

The work presented in this paper explores multiple origins of intron reads and signal in RNA-seq data. We demonstrate the usefulness of applying intron reads to study multiple aspects of transcriptional biology, and provide tools to interrogate changes in pre-mRNA and mRNA levels, as well as genes with IR-like coverage profiles. Biological validation of our results was not carried out and is beyond the scope of this paper, since the intention was to make conclusions using a data-driven approach. However, further work includes closer examination into pre-mRNA-specific and IR-specific signal, such as by using full length transcripts by long-read sequencing by Pacific Biosciences (50) or Oxford Nanopore Technologies (51). It is also of interest to examine intron reads in datasets with RNA from cytoplasmic and nuclear fractions versus whole cell.

## Conclusions

Intron reads are prevalent at small to moderate proportions in RNA-seq datasets, however, they provide signal that can distinguish between biological and experimental groups. Harvesting these extra reads as pre-mRNA signal, DGE analysis can be carried out more thoroughly with the addition of intron counts into *index*. The extra layer of analysis enables distinction between changes in pre-mRNA and mRNA signal enhancing the understanding of transcriptional changes and dynamics under study. IR remains an important mechanism in biology and its detection is possible through the use of *superintronic*, which accurately selects genes with IR-like coverage profiles, unlike that of previous methods.

## ACKNOWLEDGEMENTS

The authors would like to thank Dr Stephane Chappaz, Dr Quentin Gouil, Dr Clare Morgan and Dr Carolyn de Graaf for their helpful discussions and suggestions that have enhanced the work presented in this paper.

